# Computational design of blue melanin with peptide motif scaffolding

**DOI:** 10.64898/2026.02.02.703104

**Authors:** Di Sheng Lee, Bomi Park, Sergio Salgado, James Dolgin, David L. Kaplan

## Abstract

De novo melanin design seeks to extend natural melanin colors to new, stable colors (blue, purple, green) with sequence-to-color tunability. Natural melanin, polymerized from tyrosine (Y), is a robust pigment with heterogenous molecular weights. Control of melanin size (length) is challenging; thus, only specific colors (yellow to brown) exist in nature. In this work, we describe the design of blue melanin through the polymerization of Y-containing pentapeptides with two key properties: tight packing during peptide assembly and high solubility in aqueous environments. By motif scaffolding a pentapeptide-repeat protein (PRP) with RFdiffusion, we narrowed 160,000 possible combinations to a library of 905 Y-containing pentapeptides with tight packing features. Two of the most soluble designs successfully formed stable blue melanin with λ_max_ absorbing in 615-620 nm, contributed by homogeneous melanin length achieved around 60 Y units. Other designs also formed new colors (purple, green), along with more known colors (red, yellow, brown). We found that blue melanin exhibited thermal stability at an autoclave temperature of 121°C and photostability of weeks under 600 lux illumination. We also demonstrated the application of blue melanin as an electrophoretic ink. De novo color design from simple peptides could potentially transform how colorants are sourced and produced. Our approach with computational design should also inspire the development of new deep-learning tools to directly predict colors from amino acid sequences.

## Introduction

Blue is nature’s rarest color and therefore provides a powerful signaling cue in nature^1-4^. To appear blue, microbes and plants rely on pigmentary colorations from phycocyanins^5^ or anthocyanins^6^, while animals rely on structural coloration from nanostructured surfaces^7^. Pigmentary coloration in the animal kingdom, produced mainly by melanin, does not result in blue^8-10^, due to the heterogenous length distribution of melanin polymers and the mechanism of biosynthesis that results in broad spectral absorption. Highly conjugated macrostructures can absorb in the red (600-700 nm) region of the absorption spectrum (e.g., porphyrins^11^), albeit requiring multistep chemical reactions. The formation of long, conjugated melanin could potentially produce blue melanin in one enzymatic step, prompting the peptide design approach described in the present work to control melanin molecular weight.

Natural melanins are polymeric pigments crosslinked from tyrosine (Y) residues catalyzed by tyrosinase^12^. The first stage in the polymerization is enzyme-catalyzed, which begins with enzymatic oxidation of Ys into highly reactive quinone species^13^. The second stage of the polymerization is spontaneous, as the quinones undergo intramolecular cyclization and rearrangement for polymerization^14^. The absorption redshifts as the polymer molecular weight and length increase^15^. This system eventually polymerizes into a heterogenous mixture with wide length distribution^16^, resulting in melanin with broad spectral absorption spanning brown to black. Because of their extensive covalent crosslinking, melanin pigments are very stable towards high temperature and light exposure^17^.

Melanized peptides or melanin peptides are Y-containing peptides designed to control the heterogeneity of Y polymerization^18-20^. The peptide is designed to control the tight packing of neighboring Y residues through hydrophobic interactions and hydrogen bonds. Y residues in the peptides, upon tyrosinase oxidation, polymerize and crosslink neighboring Y-containing peptides. The resulting melanin peptides are more homogenous and have a narrower absorption peak than those from nature, thus enabling the production of pigmentary colorations of yellow, red, orange, and brown^19^. For example, pentapeptide YVPAY formed red melanin peptide with an absorption peak (λ_max_ = 430 nm) in the violet-blue region of the visible spectrum^18^.

Peptide-repeat proteins are characterized by repeating short peptide units that fold into tightly packed 3D structures^21-24^. For example, pentapeptide-repeat proteins (PRPs), with tandem repeats of five amino acids, tend to form long, compact structures^25, 26^. Each pentapeptide repeat is among the shortest known sequences capable of directing a stable tertiary structure. Four consecutive pentapeptide repeats make up one complete coil. Due to this stacking and the quadrilateral nature of the coils, the protein has four faces. For example, the crystal structure of a PRP (PDB 2BM4) is comprised of eight coils and four faces^27^. The overall structure is rod-like, with a continuous hydrophobic core of leucine or phenylalanine ladders running through the repeats and a solvent-exposed surface to maintain water solubility^28^.

Motif scaffolding is a computational protein design technique constructing de novo protein backbone (scaffold) around pre-defined protein part (motif) that matches the motif’s 3D geometry^29^. The recent development of deep learning-based RFdiffusion has substantially improved the accuracy of motif scaffolding^30, 31^. For example, this approach has been used to perform various motif scaffoldings to design de novo enzymes^32, 33^ and antivenom proteins^34^.

The design of blue melanin from Y-containing pentapeptides with 4 variable residues requires a reduction from 160,000 (20^4^) possible combinations of the amino acids. Here, we demonstrate the design of blue melanin from Y-containing pentapeptides by motif scaffolding a PRP (Fig. 1a-c). First, RFdiffusion was used to generate a library of Y-containing pentapeptides with tight packing features that can form a Y ladder. Second, the library of peptides was subjected to a solubility screen to select for the most soluble Y-containing pentapeptides. This led to the selection of 10 motif scaffolding designs maximized for solubility for experimental validation. We demonstrated that 2 designs successfully formed blue melanin: SPYGG and LAYPS, with a narrow absorption peak in the red region of the absorption spectra.

**Fig. 1:**
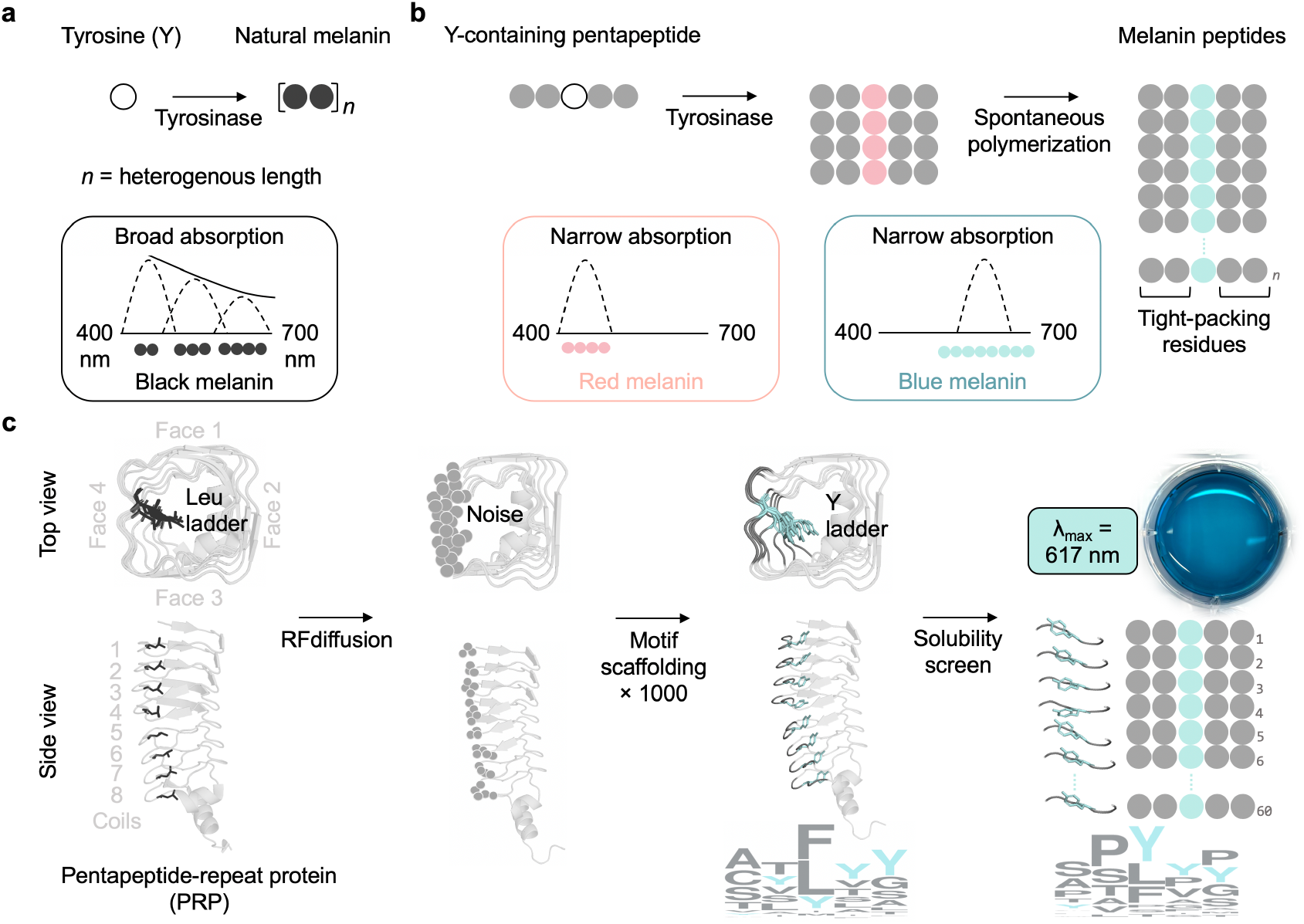
Computational design of blue melanin. **a**, The formation of natural melanin polymer from Y residues. After tyrosinase oxidation, the Y residues are crosslinked through covalent bonds. The heterogeneous mixture of short and long melanin polymers produces a broad absorption spectrum that gives melanin its black color. **b**, The formation of melanin peptides (or melanized peptides) from Y-containing pentapeptides. The first stage of melanin polymerization is tyrosinase-catalyzed. The second stage of the polymerization is spontaneous. The homogenous length distribution of melanin peptides produces narrow absorption spectra that give melanin peptides red or blue colors. Melanin peptide length, *n*, is tunable via tight-packing residues. **c**, Sequence design of tight-packing residues with RFdiffusion. A PRP (PDB 2BM4) is motif scaffolded 1000 times on face 4 to generate scaffold designs of tightly packed Y-containing pentapeptides. After solubility screen, the sequence design of the top 20 peptides was shown. An absorption peak with a λ_max_ of 617 nm is produced by long blue melanin peptides with *n* = 60, crosslinked by Y units located in the middle of the pentapeptide.

## Results

### Motif scaffolding of PRP

Y-containing pentapeptides that tightly pack should produce long melanin polymers that have blue pigmentary colorations. Ideally, they have a single absorption peak in the red region of the visible spectrum. Here, we used RFdiffusion to design Y-containing pentapeptides with tight-packing features by motif scaffolding a PRP (PDB 2BM4). The PRP was partially scaffolded to replace native pentapeptide repeats with Y-containing sequences that can form a Y ladder (Fig. 1c). Faces 1-3 of PRP were used as motifs, and face 4 was scaffolded to custom generate Y-containing pentapeptides (Methods). The role of face 1-3 motifs was to keep the protein structure in the exact same 3D quadrilateral geometry with 4 faces and 8 coils. We input the 3D coordinates of faces 1-3, and random noise was added to face 4. RFdiffusion iteratively refined that noise until new backbones were generated through conditioned denoising trajectories.

Following the generation of new backbones on face 4, 1,000 sequence designs were carried out using ProteinMPNN to generate Y-containing sequences. We avoid sampling these 8 residues (R, H, K, D, E, W, N, Q) because charged, highly polar, and bulky residues compromise compact packing^18^. To increase the diversity of the designs for screening, a high sampling temperature of 1.0 was applied to the remaining 12 residues (A, C, F, G, I, L, M, P, S, T, V, Y). Since each PRP has 8 coils, a total of 8,000 pentapeptide designs with high confidence (>0.85 pLDDT and >0.8 pTM scores) were generated from the resulting 1,000 PRPs (Supplementary Fig. 1). Out of the 8,000 pentapeptides, there were 905 Y-containing pentapeptides: 855 peptides with 1 Y and 50 peptides with 2 Ys (Fig. 2a).

**Fig 2:**
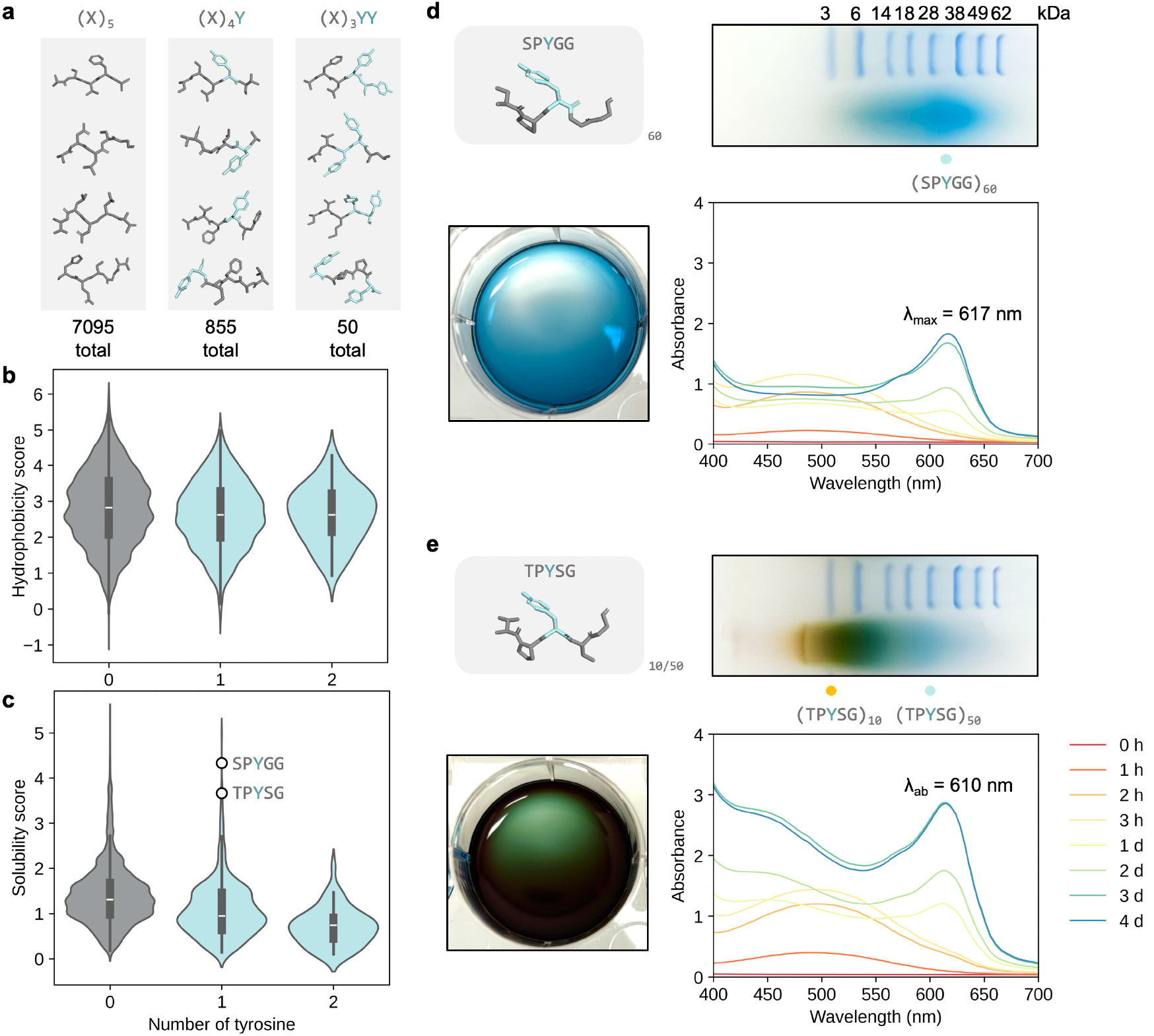
Experimental characterization of top Y-containing pentapeptides. **a**, Scaffold library of 8,000 motif scaffolded pentapeptides with 0, 1, or 2 Ys. **b**, Hydrophobicity distribution of 8,000 motif scaffolded peptides. **c**, Solubility distribution of 8,000 motif scaffolded peptides. **d**, Absorption spectrum and native PAGE gel of blue melanin formed from SPYGG peptide with a λ_max_ of 617 nm. **e**, Absorption spectrum and native PAGE gel of green melanin formed from TPYSG peptide.

High solubility should increase the maximum concentration a peptide can reach in water, which is important for tight packing^35^. To achieve high solubility, all 8,000 pentapeptides were analyzed and scored for hydrophobicity and solubility (Methods) (Fig. 2b, c). The residue frequency and diversity of the peptide designs before and after solubility screen were analyzed with sequence logo method (Fig. 1c). Ten Y-containing pentapeptide designs were selected for experimental validation based on ranking of their solubility scores (Supplementary Fig. 2 and Supplementary Table 1).

### Production of blue melanin peptides

The best Y-containing peptides designed by motif scaffolding were those that had high solubility in water. The peptides were chemically synthesized with purity of >98% to test for their ability to form blue melanin after tyrosinase oxidation. As controls, we also selected three peptides with lower solubility (TSYAG, STYLM, and CTYFL) and the top 3 peptides with 2 Ys (SYYGG, YSYVG, and SYMYG) (Supplementary Fig. 2 and Supplementary Table 2). The peptides were resuspended in 0.05 M sodium phosphate buffer at pH 7 at a concentration of 10 mg/mL (Methods). Mushroom tyrosinase from *Agaricus bisporus* was added to initiate the first stage of enzymatic oxidation for 3 hours (Methods). The oxidative polymerization of Ys in the pentapeptide was confirmed by observing a gradual change in color and measuring the formation of the absorption peak in the visible spectrum (400-700 nm) (Methods). After 3 hours of oxidation, the peptide solutions were incubated at −20°C to support spontaneous polymerization with reduced conformational movement in order to facilitate enhanced tight packing of pentapeptides. Here the goal was to maximize melanin polymer length to absorb red wavelengths. The gradual red shift of the absorption peak continued for at least 4 days and up to 10 days.

The second most soluble peptide, SPYGG successfully formed blue melanin, with an absorption peak at 617 nm after 4 days of polymerization (Fig. 2d). The solution turned dark red after 3 hours of oxidative polymerization at room temperature (Supplementary Fig. 2). On day 2, the frozen solution turned to gray-black. On day 4, blue melanin formed with a λ_max_ of 617 nm with a narrow bandwidth of 60 nm. The second stage of polymerization was spontaneous, so the reaction proceeded even in frozen form at −20°C. In contrast, spontaneous polymerization at room temperature did not result in blue melanin formation (Supplementary Fig. 2). Native polyacrylamide gel electrophoresis (PAGE) gel showed that the blue melanin had a length of 10-100 Y units, with the most intense blue band at 60 Y units.

The third most soluble peptide, TPYSG formed green melanin, with an absorption peak at 610 nm and strong 400-500 nm absorption (Fig. 2e). Native PAGE gel showed that green melanin consisted of a combination of yellow and blue melanins, which had chain lengths of 10 and 50 Y units, respectively. The yellow melanin was responsible for the strong absorption in the 400-500 nm, and the blue melanin was responsible for the strong absorption at 610 nm. Thus, TPYSG was a less stable derivative of blue melanin.

The absorbance spectra of the remaining 8 of the top 10 peptides (PPYVA, VSYPG, SSFPY, SPLYG, CAPYS, YPFAS, LAYPS, and ASLYP) and low-solubility peptides (TSYAG, STYLM, and CTYFL) were obtained (Fig. 3). PPYVA underwent rapid polymerization and formed purple melanin with an absorption peak at 530 nm in the first stage of polymerization. The peak did not red shift even after 10 days of spontaneous polymerization at −20°C. VSYPG formed red melanin, while ASLYP, YPFAS, and STYLM formed brown melanins. Absorption peaks in the 600-700 nm were present in SPLYG, SSFPY, LAYPS, and TSYAG melanins, suggesting partial blue melanin formations. After 10 days, LAYPS successfully formed blue melanin with a λ_max_ of 619 nm. Cysteine-containing peptides (CTYFL and CAPYS) did not polymerize, consistent with previous studies that cysteine inhibits melanin formation^36, 37^ (Fig. 3 and Supplementary Fig. 2).

**Fig 3:**
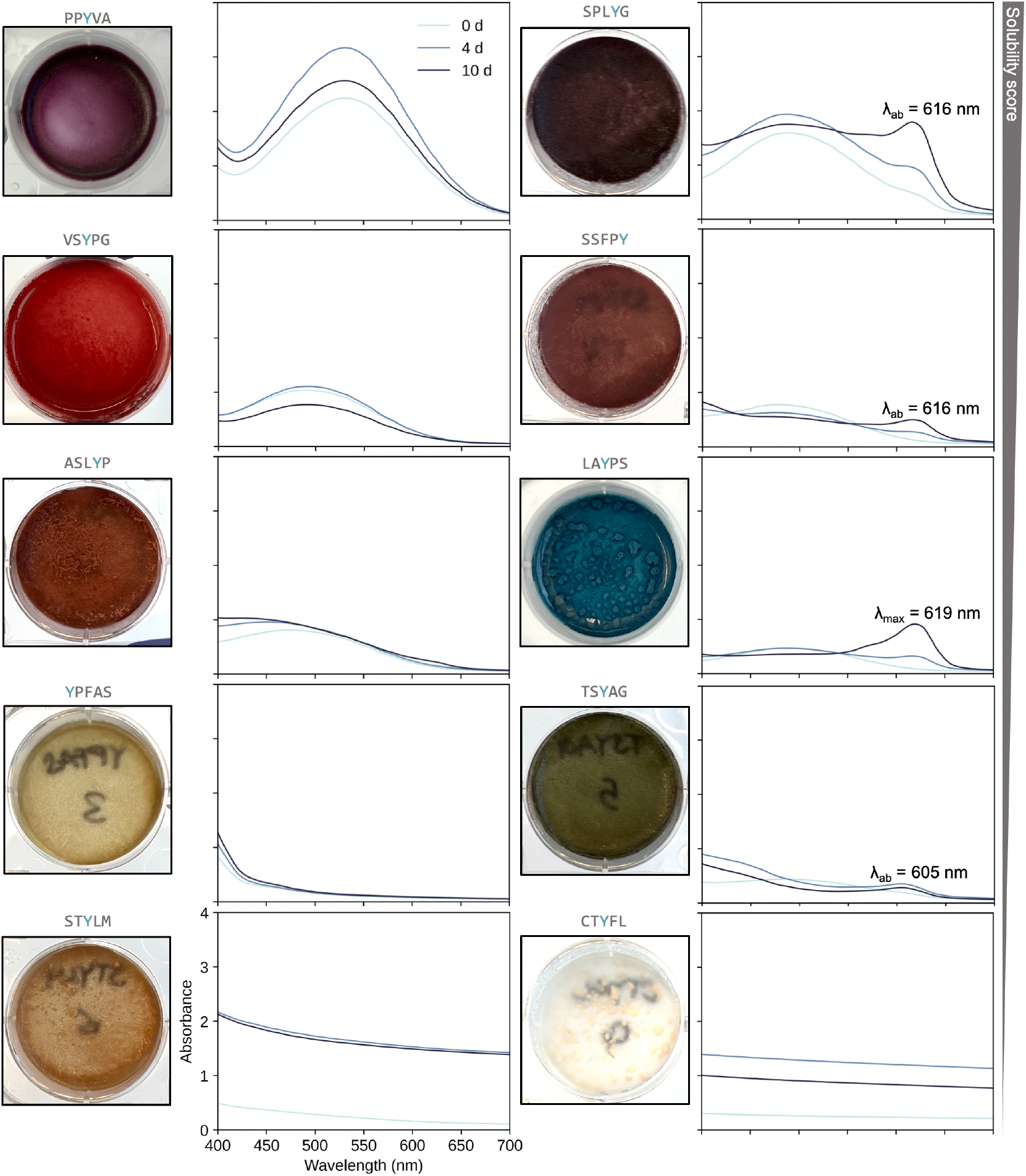
Formation of melanin peptides with various pigmentary colorations. Absorption spectra of melanin peptides formed from Y-containing pentapeptides. Absorption peaks in the red region of the visible spectrum are present in SPLYG, SSFPY, LAYPS, and TSYAG melanins. The solubility scores of the peptides are 5.0 (PPYVA), 3.7 (SPLYG, VSYPG, SSFPY), 3.4 (ASLYP, LAYPS, YPFAS), 1.9 (TSYAG), 1.0 (STYLM), and 0.3 (CTYFL). The solubility ranking is provided in Supplementary Fig. 2 and Supplementary Table 1.

The presence of 2 Ys in the pentapeptide sequence reduced the solubility and did not lead to blue melanin formation. By substituting proline in SPYGG with another Y into SYYGG, the solubility decreased significantly, resulting in precipitation of the peptide upon resuspension in 0.05 M sodium phosphate buffer (Supplementary Fig. 2). SYMYG with low solubility precipitated in 0.05 M sodium phosphate buffer. While YSYVG was soluble, it had a red-brown appearance.

The two blue melanin precursors (SPYGG and LAYPS) were selected to produce hybrid blue melanin. We mixed SPYGG and LAYPS peptides at equal weights and polymerized the mixture for a total of 10 days. On day 4, strong absorption in 400-600 nm was still present (Supplementary Fig. 3). On day 10, the shorter peptides previously absorbing in the 400-600 nm range polymerized into longer melanins, producing a hybrid blue melanin with a λ_max_ of 620 nm.

Oxygen availability is critical to melanin formation^38^. The use of a petri dish was important to ensure high surface area to volume ratio for oxygen exchange with the peptide solution. Using a microcentrifuge tube for cold polymerization significantly delayed SPYGG blue melanin formation to 2 weeks (Supplementary Fig. 4). Similarly, despite the −20°C, this stage of polymerization was spontaneous and therefore was able to proceed in frozen form. However, with limited oxygen supply, the spontaneous polymerization rate was much slower.

Evaporation has the same effect of maximizing tight packing of peptides by concentrating the peptides as the water evaporates. The reactions were carried out in petri dish without a lid to allow for evaporation at room temperature. After 1 day of polymerization and evaporation, the evaporated solution turned blue (Supplementary Fig. 4). However, this technique was less controllable and the final color after full evaporation was green.

### Molecular dynamics of low-temperature tight packing

Peptide clusters should have minimal thermal motions at low temperatures. Ideally, the peptide clusters should have minimal change in solvent accessible surface area (SASA) at −20°C, thus facilitating tight packing, bond formation, and polymerization. Atomistic molecular dynamics simulations are lengthy and computationally intensive^39^. To simulate the tight packing biophysics of SPYGG peptides at different temperatures in the milliseconds, coarse-grained molecular dynamics were carried out^40^.

The SPYGG peptide was converted to a coarse-grained representation using MARTINI force field v2.6 (Methods). The simulation consisted of 462 coarse-grained SPYGG peptides randomly placed in a box with a size of 15 × 15 × 15 nm (Supplementary Fig. 5). Water was explicitly represented as beads with 0.21 nm van der Waals radii. SPYGG peptides were allowed to pack into a cluster at 30°C until the SASA stabilized, which occurred around 0.55 ms. After the cluster stabilized, the simulations were allowed to proceed for another 0.15 ms to study the fluctuation of SASA between 30°C and −20°C. The effect of temperature on tight packing could be inferred from SASA fluctuation from 0.55-0.70 ms: low temperature minimizes thermal movements and maximizes tight packing.

### Stability of SPYGG blue melanin

Once SPYGG blue melanin was formed after at least 4 days of polymerization, the pH-, shelf-, light-, and thermal stabilities of SPYGG blue melanin were evaluated. For each stability test, SPYGG blue melanin was re-suspended to a final absorbance value of 0.55. If the color was not stable under the prescribed condition (e.g., its absorbance peak decreased), the pH was adjusted from pH 7 to pH 5, 3, and 1 until stability was observed.

First, we tested the stability of SPYGG blue melanin across pH 1-13. The color change and change in absorbance peak were observed (Fig. 4a). An increase in absorbance in acidic conditions was observed, suggesting that low pH facilitates blue melanin formation. The red shift and decrease in absorbance in basic conditions were consistent with depolymerization and alkaline hydrolysis of melanin^41^. We compared the pH stability against commercial synthetic blue dyes such as Blue 1 and Blue 2 (Supplementary Fig. 6). We also compared the blue melanin against commercial natural dyes such as spirulina phycocyanin and blueberry anthocyanin. The relative stability of SPYGG blue melanin was found to be between those of synthetic and natural blue dyes.

**Fig 4:**
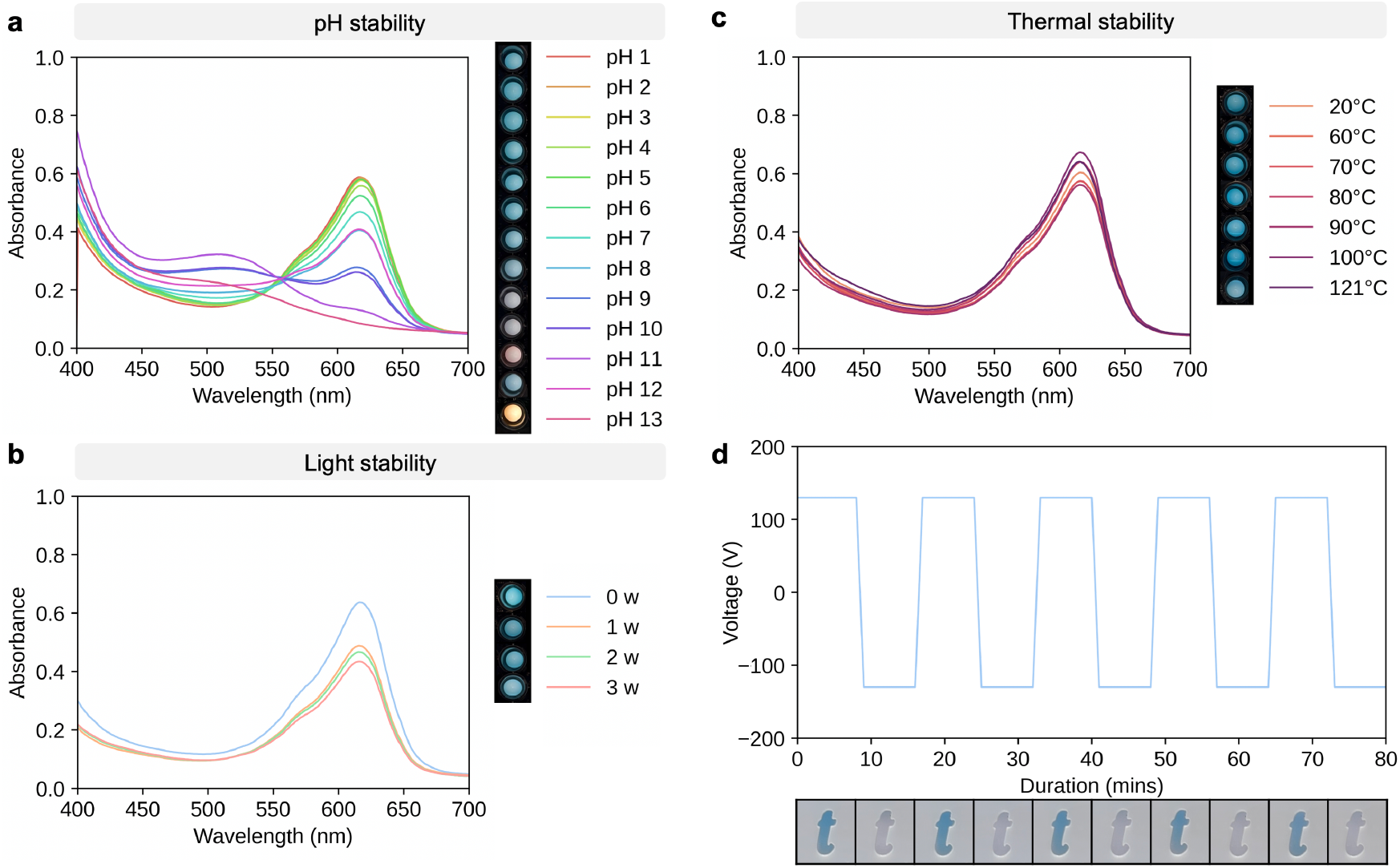
Stabilities of SPYGG blue melanin. **a**, Absorption spectra of SPYGG blue melanin re-suspended in pH 1-13 buffer solutions. Each spectrum is average of three replicates. **b**, Absorption spectra of SPYGG blue melanin re-suspended in pH 1 buffer solution exposed to 600 lux of illumination over 3 weeks. Each spectrum is average of three replicates. **c**, Absorption spectra of SPYGG blue melanin re-suspended in pH 1 buffer solution heated to 60°C, 70°C, 80°C, 90°C, 100°C and 121°C (autoclave) for 30 mins. Each spectrum is average of three replicates. **d**, Application of SPYGG blue melanin as electronic ink under an electric field of 130 V with polarity reversing every 8 mins for a total of 80 mins.

Next, we tested the shelf stability of SPYGG blue melanin stored in the dark for 3 weeks. At neutral pH, the absorbance peak decreased with time, suggesting gradual depolymerization of blue melanin (Supplementary Fig. 7). To protect from depolymerization, the SPYGG blue melanin was resuspended in a pH 5 buffer solution and stored in the dark for 3 weeks. No significant drop in absorbance peak was observed, suggesting that it is shelf-stable at pH 5. Similar stability at pH 5 was observed in TPYSG green melanin (Supplementary Fig. 7).

We then tested the light stability of SPYGG blue melanin under illumination of 600 lux over 3 weeks. At neutral pH, a significant drop in absorbance peak was observed in the first week, suggesting rapid photodegradation and depolymerization of blue melanin (Supplementary Fig. 8). To improve its stability against photodegradation, SPYGG blue melanin was resuspended in pH 5, 3, and 1 buffer solutions, respectively. No significant drop in the absorbance peak was observed at pH 1, indicating photostability at this pH (Fig. 4b). We compared this light stability against commercial synthetic and natural blue (Supplementary Fig. 9). The light stability of SPYGG blue melanin was between those of synthetic and natural blue dyes.

Lastly, we tested the thermal stability of SPYGG blue melanin at 60°C, 70°C, 80°C, 90°C, 100°C and 121°C (autoclave) for 30 mins. At neutral pH, significant decreases in absorbance were observed at all elevated temperatures (Supplementary Fig. 8). To improve stability against thermal degradation, SPYGG blue melanin was resuspended in pH 5, 3, and 1 buffer solutions, respectively. At pH 5 and 3, SPYGG blue melanin was stable from 60°C to 100°C. At pH 1, SPYGG blue melanin was stable even after autoclaving temperature of 121°C (Fig. 4c). We also compared the thermal stability against commercial synthetic and natural blue (Supplementary Fig. 10). SPYGG blue melanin was thermally more stable than the natural blue dyes, but not the synthetic ones.

### Application of blue melanin as electrophoretic ink

Natural melanin could be used as electronic ink that migrates under an electric field for electrophoretic display^42^. An electrophoretic display was prepared by overlaying a mask consisted of the letter “t” on top of the running gel apparatus (Methods). A running buffer with neutral pH was used since SPYGG blue melanin was not stable in a common running buffer of pH 8.4 (Methods). An electric field of 130 V was applied, and the direction of the electric field was reversed every 8 mins for a total of 10 cycles. Because melanin is intrinsically negatively charged due to its −COO− groups, the peptide migrated towards the positively charged anode. A blue letter t could be observed through the naked eye when a positive electric field was applied (Fig. 4d).

## Discussion

Natural melanin, prepared from Y, is one of the most stable natural pigments. It can withstand high temperatures beyond boiling and does not photodegrade under exposure of ultraviolet light. While these natural melanins can be found in brown-black colors in the form of eumelanin, or red-yellow color in the form of pheomelanin, the formation of blue melanin remains a challenge^43, 44^. Here, we introduce design principles of melanin peptides that, when combined with deep learning-based protein design tools such as RFdiffusion, enable the preparation of long melanin polymers that display blue coloration. From peptide sequence to color, we have successfully designed de novo melanin colors (blue, purple, and green) and known colors (red, yellow, brown) using this approach. Further, we showed that blue melanin survived autoclaving temperature of 121°C, did not photodegrade after weeks of light exposure, and can be used as an electrophoretic ink.

This work involved computational design principles to control the intrinsic heterogenous polymerization of melanin from Y residues. The designs that we explored are derived from Y-containing pentapeptides. Tight-packing motif scaffolding designs, followed by solubility screens, lead to melanin peptides with single absorption peak in the red region of the visible spectrum. Melanin starts absorbing light after tyrosinase oxidation, due to the formation of melanin oligomers with highly conjugated backbones. As the length of conjugated melanin increases, the absorption peak red shifts to longer wavelengths, thus allowing absorption further into the red region than eumelanin or pheomelanin. The single absorption peak or narrow absorption spectrum is due to the formation of homogeneous melanins with narrow (more homogenous) length distribution. Tightly packed, highly soluble peptides align to bring the Y residues together to form homogeneous melanin polymers of controllable lengths, providing a lever for sequence-to-color tuning to create de novo blue, purple, and green melanins.

It is widely believed that blue colorations observed in bird feathers are due to structural color^45-47^. We demonstrate that melanin derived from Y can produce blue pigmentary colorations, due to the formation of long melanin polymers that can be as long as 100 Ys. We observed that the formation of blue melanin was enhanced in acidic conditions. This is consistent with previous studies that the formation of uniform, very high molecular weight melano–protein complexes was enhanced in acidic organelles called melanosomes^48, 49^. The low pH environment maximizes the interaction between protein and melanin^50^. Our work suggests that blue feathers could also be due to pigmentary colorations after all, and not just structural color. Further, the acidic intra-melanosomal milieu could play a key role in forming and stabilizing these blue melanins, preventing these high molecular weight melano-protein complexes from photobleaching.

Although only melanin peptides were experimentally tested in this work, the potential of melanin proteins such as SPYGG-repeat proteins could be much bigger. One can imagine the application of motif scaffolding to create blue melanin proteins that are much more stable and homogeneous. Peptide-repeat proteins are attractive targets for melanin protein design and prime structural motifs for RFdiffusion^51^. The ability to customize RFdiffusion to specific design challenges by the addition of proper moieties and by fine-tuning amino acid diversity in the scaffold sequence should enable higher levels of complexity and more specific designs. The tandem repeat design of peptide-repeat proteins represents a simple correspondence between amino acid sequence and Y polymerization length, enabling tunable control over the length and hence color of Y-based melanin.

Deciphering peptide-based colors could unlock new computational tools that predict color solely from peptide sequence of various lengths. The melanin peptides we designed are 5-residue long; however, shorter or longer melanin peptides can also be designed using tetrapeptide-, hexapeptide-, or heptapeptide-repeat motifs. Although we have focused on chemically synthesized peptides, these melanin peptides could potentially be formed in vivo through the co-expression of peptides and tyrosinase in acidic organelle similar to melanosome for imaging applications. Further, such strategies could include domains for anchoring techniques, to further the assembly and stability of the pigments that form. By enabling de novo color design from sequence to color in nature’s fourth biopolymer^52^, melanin peptide-based colorations could revolutionize how colorants are sourced and produced in the food dyes, cosmetics, and paints industries. Since these are peptide-based solutions, the pigments should be human and environmentally compatible.

## Materials and Methods

### Materials

Peptides were chemically synthesized and purchased from Genscript (Piscataway, NJ, U.S.A.) at 98% purity followed by high-performance liquid chromatography. We dissolved the lyophilized peptide at room temperature in Ultrapure water (Invitrogen, 10977015) to a final concentration of 10 mg/mL. Tyrosinase (Sigma #T3824) was dissolved to 2 mg/mL in Ultrapure water. 0.1 M sodium phosphate buffer, pH 7 was prepared from sodium dihydrogen phosphate dihydrate (Sigma #1063420250) and sodium phosphate dibasic dodecahydrate (Sigma #71663). Colorless pH 1, 2, 3, 5, 6, 8, 9, 11, 12, and 13 buffer solutions (ThermoFisher #040446, #040447, #042416, #042417, #042418, #042420, #042421, #042424, #042425, #046307) were used for re-suspending melanin peptides. Colorless pH 4 and pH 10 were prepared by adjusting with hydrochloric acid (Boston BioProducts #BZ-8010) and sodium hydroxide solution (Sigma #S2770). Brilliant Blue FCF (Sigma #80717) was used as synthetic blue dye No. 1. Indigo Carmine (ThermoFisher #412300250) was used as synthetic blue dye No. 2. USDA organic blue spirulina phycocyanin extract powder (Asiya Life Company Limited #B08YRDBFFX) was used as natural phycocyanin dye. Organic blueberry powder (Bulk Supplements #BLUPWD500) was used as natural anthocyanin dye.

### RFdiffusion motif scaffolding

The crystal structure of PRP (2BM4) was used as inputs for RFdiffusion v1.1.1 on Google Colab. Faces 1-3 of the PRP were used as structural motifs, and face 4 was scaffolded to custom generate Y-containing pentapeptides: contigs=‘A1-16/5/A22-36/5/A42-56/5/A62-76/5/A82-96/5/A102-116/5/A122-136/5/A142-156/5/A162-183’; iterations=‘50’; num_designs=‘1’. 1,000 diffused designs were generated. Sequence design was then carried out on the backbone libraries using ProteinMPNN, followed by FastRelax and AF2 + initial guess^53^. Charged, highly-polar, and bulky residues (R, H, K, D, E, W, N, Q) were not sampled. A high sampling temperature of 1.0 was apply to the remaining residues (A, C, F, G, I, L, M, P, S, T, V, Y). The resulting libraries were filtered using predefined filters: AF2 predicted aligned error (PAE) < 10 and predicted local distance difference test (pLDDT) > 0.85.

### Coarse Grained Molecular Dynamics

The protocol was modified from previous work^18, 54^. The coordinate files for SPYGG peptide was created and coarse-grained using MARTINI force field v2.6^55^ and GROMACS v2019.2^56^. A cubic box of 15nm × 15nm × 15nm was used. 462 pentapeptides were randomly inserted. Standard Martini coarse-grained water was used. The periodic boundary conditions, Lennard-Jones interactions, and electrostatic interactions were kept the same as previous work^54^. The system was first energy-minimized for 300,000 steps to lower maximum force on each atom to below 200 pN. The system was allowed to equilibrate for 22,00,000 steps using a 25 fs timestep (0.55 ms). The temperature and pressure were 303 K and 1 bar during the equilibrium. The system was further allowed to equilibrate at 303 K and 253 K respectively for 6,000,000 steps using a 25 fs timestep (0.15 ms), resulting in a total simulation time of 0.70 ms.

### Surface area, hydrophobicity, and solubility calculation

SASA was calculated with *gmx sasa* command in GROMACS v2019.2. A probe size of 0.14 nm (similar to water molecule size) was used to determine how much of the peptide cluster was accessible by the probe. Hydrophobicity score of each peptide was calculated as

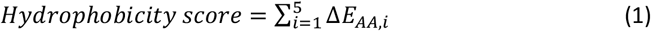

where Δ*E*_*AA,i*_is the Eisenberg value of each amino acid^57^.

The solubility score was calculated as

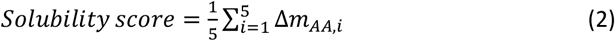

where Δ*m*_*AA,i*_is the molality of each amino acid at 25°C, obtained from previous study^58^.

### Melanin peptide polymerization

The peptide was dissolved in 2 mL of 0.05 M sodium phosphate buffer at room temperature (20°C) to a final concentration of 10 mg/mL in a 35 mm diameter petri dish (Falcon #351008). The solution was allowed to equilibrate for 5 min. Tyrosinase solution was then added to the peptide solution to a final concentration of 0.2 mg/mL. The oxidation reaction was performed with lid in two stages. The first stage of enzymatic polymerization was allowed to proceed at room temperature for 3 hours. The second stage of spontaneous polymerization was carried out at −20°C for at least 4 days.

### Measurement of absorption spectra

A 60 µL aliquot of melanin peptide was added in a 96-well black polystyrene microplate (Corning #3603). Absorption spectra of melanin peptides were measured in the 400-700 nm range with interval of 1 nm using BioTek Synergy H1 plate reader (Biotek # 11-120-533). The measurements were performed at room temperature (20°C).

### Time course of melanin peptide polymerization

In the first stage of enzymatic polymerization, 60 µL of melanin peptide was transferred from petri dish to a 96-well black microplate for absorbance measurement after 1, 2, and 3 hours of oxidation. In the second stage of spontaneous polymerization, the frozen solution was partially thawed at room temperature. A 60 µL aliquot of melanin peptide was transferred from the petri dish to a 96-well black microplate for absorbance measurement after 1, 2, 3, and 4 days of oxidation (up to 10 days).

### Native PAGE

Native PAGE was performed using 12% pH 6.4 Bis-Tris gels (Invitrogen #NW00120BOX) in mini gel tank (Invitrogen #A25977) powered by Consort EV 265 electrophoresis power supply (ReScience #2411270004). Then 20 µL of SPYGG blue melanin or TPYSG green melanin was mixed with 2 ul of glycerol (Sigma #G5516). Samples were loaded into the gel in MES running buffer (Invitrogen #B0002) and run at 130 V for 30 mins. SeeBlue™ Pre-stained Protein Standard ladder (ThermoFisher #LC5625) was used. Once finished, gels were rinsed with water and imaged with a cell phone on a ChemiDoc™ Imaging System white sample tray (Bio-Rad #12003026).

### Electrophoretic display

A mask consisted of the letter “t” (Amazon #B0FK4N5XD8) was placed on top of the running gel apparatus. 20 µL of SPYGG blue melanin was mixed with 2 ul of glycerol. MES running buffer with neutral pH was used. An electric field of 130 V was applied by Consort EV 265 electrophoresis power supply. The electric field was reserved by reversing the black and red cables every 8 mins for a total of 10 cycles. The t-letter mask was periodically imaged every 8 mins.

### pH stability

After at least 4 days of spontaneous polymerization at −20°C, SPYGG blue melanin was re-suspended in colorless pH1-13 buffer solutions to a final absorbance value of 0.55. The re-suspended solution was allowed to equilibrate for 30 mins. Next, 60 µL aliquot of the solution was transferred to a 96-well black microplate for absorbance measurement. Synthetic and natural pigments were prepared using the same protocol.

### Shelf stability

After at least 4 days of spontaneous polymerization at −20°C, SPYGG blue melanin and TPYSG green melanin were re-suspended in colorless pH 5 and 7 buffer solutions to a final absorbance value of 0.55 in 1.5 mL microcentrifuge tubes (USA Scientific #1415-2500). The sample was kept in dark at room temperature. 60 µL aliquot of melanin peptide was transferred to a 96-well black microplate for absorbance measurement after 0, 1, 2, and 3 weeks.

### Light stability

After at least 4 days of spontaneous polymerization at −20°C, SPYGG blue melanin was re-suspended in colorless pH 1, 3, 5, and 7 buffer solutions to a final absorbance value of 0.55 in 1.5 mL microcentrifuge tubes. Similarly, synthetic Blue 1, Blue 2, and spirulina phycocyanin were re-suspended in colorless pH 7 buffer solutions. Blueberry anthocyanin was re-suspended in colorless pH 11 buffer. The samples were exposed to light source (Lepro #PR310001-DWW) under 600 lux of illumination measured by a light meter (URCERI, #SMT912). Then 60 µL aliquot of melanin peptide was transferred to a 96-well black microplate for absorbance measurement after 0, 1, 2, and 3 weeks of continuous light exposure.

### Thermal stability

After at least 4 days of spontaneous polymerization at −20°C, SPYGG blue melanin was re-suspended in colorless pH 1, 3, 5, and 7 buffer solutions to a final absorbance value of 0.55 in 1.5 mL microcentrifuge tubes. Similarly, synthetic Blue 1, Blue 2, and spirulina phycocyanin were re-suspended in colorless pH 7 buffer solutions. Blueberry anthocyanin was re-suspended in colorless pH 11 buffer. The samples were heated at 60°C, 70°C, 80°C, 90°C, and 100°C in a block heater (ThermoFisher #88871001) for 30 mins. For thermal stability above 100°, the samples were autoclaved for 30 mins. Next, 60 µL aliquot of melanin peptide was transferred to a 96-well black microplate for absorbance measurement.

## Acknowledgements

This research was sponsored by the United States Department of Agriculture and was accomplished under Cooperative Agreement Number MASW-2021-05678. The views and conclusions contained in this document are those of the authors and should not be interpreted as representing the official policies, either expressed or implied, of the United States Department of Agriculture or the U.S. Government. The U.S. Government is authorized to reproduce and distribute reprints for Government purposes notwithstanding any copyright notation herein.

## Author information

### Authors and Affiliations

Department of Biomedical Engineering, Tufts University, Medford, MA 02155, USA

Di Sheng Lee, Bomi Park, Sergio Salgado, James Dolgin, and David L. Kaplan

Tufts University Center for Cellular Agriculture, Medford, MA 02155, USA

Di Sheng Lee, Bomi Park, Sergio Salgado, James Dolgin, and David L. Kaplan

### Contributions

D.S.L. and D.L.K. conceptualized the project, designed experiments, and wrote the paper. D.S.L. designed, screened, and experimentally characterized all melanin peptides shown in this paper. B.P. assisted with stability tests on synthetic and natural dyes. S.S. and J.D. assisted with melanin peptide size characterization. All authors reviewed and accepted the paper.

## Competing interests

The authors declare no competing interests.

## Data availability

Peptide sequence data used in this work are included in Supplementary Information. Source data will be provided with this paper. These and any additional data are available from the corresponding author upon request.

## Notes

### Competing Interest Statement

The authors have declared no competing interest.

## References

1. Berke, H. The invention of blue and purple pigments in ancient times. Chem Soc Rev 36, 15–30 (2007).

2. Kupferschmidt, K. In search of blue. Science 364, 424–429 (2019).

3. Newsome, A.G., Culver, C.A. & van Breemen, R.B. Nature’s palette: the search for natural blue colorants. J Agric Food Chem 62, 6498–6511 (2014).

4. Umbers, D.L. On the perception, production and function of blue colouration in animals. Journal of Zoology, 229–242 (2012).

5. Eriksen, N.T. Production of phycocyanin--a pigment with applications in biology, biotechnology, foods and medicine. Appl Microbiol Biotechnol 80, 1–14 (2008).

6. Santos-Buelga, C., Mateus, N. & De Freitas, V. Anthocyanins. Plant pigments and beyond. J Agric Food Chem 62, 6879–6884 (2014).

7. Bagnara, J.T., Fernandez, P.J. & Fujii, R. On the blue coloration of vertebrates. Pigment Cell Res 20, 14–26 (2007).

8. Riley, P.A. Melanin. Int J Biochem Cell Biol 29, 1235–1239 (1997).

9. Ito, S. & Wakamatsu, K. Quantitative analysis of eumelanin and pheomelanin in humans, mice, and other animals: a comparative review. Pigment Cell Res 16, 523–531 (2003).

10. McGraw, K.J. An update on the honesty of melanin-based color signals in birds. Pigment Cell Melanoma Res 21, 133–138 (2008).

11. Chemla, Y. et al. Hyperspectral reporters for long-distance and wide-area detection of gene expression in living bacteria. Nat Biotechnol (2025).

12. Sanchez-Ferrer, A., Rodriguez-Lopez, J.N., Garcia-Canovas, F. & Garcia-Carmona, F. Tyrosinase: a comprehensive review of its mechanism. Biochim Biophys Acta 1247, 1–11 (1995).

13. Land, E.J., Ramsden, C.A. & Riley, P.A. Quinone chemistry and melanogenesis. Methods Enzymol 378, 88–109 (2004).

14. Mondal, S., Thampi, A. & Puranik, M. Kinetics of Melanin Polymerization during Enzymatic and Nonenzymatic Oxidation. J Phys Chem B 122, 2047–2063 (2018).

15. Arzillo, M. et al. Eumelanin buildup on the nanoscale: aggregate growth/assembly and visible absorption development in biomimetic 5,6-dihydroxyindole polymerization. Biomacromolecules 13, 2379–2390 (2012).

16. Choudhury, A. & Ghosh, D. Elucidating the Structure of Melanin and Its Structure-Property Correlation. Acc Chem Res 58, 1509–1518 (2025).

17. Xie, W. et al. Natural Eumelanin and Its Derivatives as Multifunctional Materials for Bioinspired Applications: A Review. Biomacromolecules 20, 4312–4331 (2019).

18. Lee, D.S., Yuan, W. & Voigt, C.A. Infrared-Scattering by Programmable Peptide-Melanin Microparticles. Advanced Functional Materials 34 (2024).

19. Lampel, A. et al. Melanin-Inspired Chromophoric Microparticles Composed of Polymeric Peptide Pigments. Angew Chem Int Ed Engl 60, 7564–7569 (2021).

20. Lampel, A. et al. Polymeric peptide pigments with sequence-encoded properties. Science 356, 1064–1068 (2017).

21. Zeytuni, N. & Zarivach, R. Structural and functional discussion of the tetra-trico-peptide repeat, a protein interaction module. Structure 20, 397–405 (2012).

22. Pretorius, D. et al. Designing novel solenoid proteins with in silico evolution. Commun Chem 9, 10 (2025).

23. Nikov, G.I., Pretorius, D. & Murray, J.W. SOLeNNoID: a deep learning pipeline for solenoid residue detection in protein structures. Bioinformatics 41 (2025).

24. Arrías, P.N. et al. Diversity and structural-functional insights of alpha-solenoid proteins. Protein Science 33, e5189 (2024).

25. Bateman, A., Murzin, A.G. & Teichmann, S.A. Structure and distribution of pentapeptide repeats in bacteria. Protein Sci 7, 1477–1480 (1998).

26. Vetting, M.W. et al. Pentapeptide repeat proteins. Biochemistry 45, 1–10 (2006).

27. Hegde, S.S. et al. A fluoroquinolone resistance protein from Mycobacterium tuberculosis that mimics DNA. Science 308, 1480–1483 (2005).

28. Calia, C.N. & Paesani, F. Computational Analysis of Threonine Ladders on Distinct beta-Solenoid Scaffolds, with Implications for the Design of Novel Antifreeze Proteins. J Phys Chem B 129, 9357–9372 (2025).

29. Wang, J. et al. Scaffolding protein functional sites using deep learning. Science 377, 387–394 (2022).

30. Watson, J.L. et al. De novo design of protein structure and function with RFdiffusion. Nature 620, 1089–1100 (2023).

31. Ahern, W. et al. Atom-level enzyme active site scaffolding using RFdiffusion2. Nat Methods 23, 96–105 (2026).

32. Braun, M. et al. Computational enzyme design by catalytic motif scaffolding. Nature 649, 237–245 (2026).

33. Kim, D. et al. Computational design of metallohydrolases. Nature 649, 246–253 (2026).

34. Torres, S.V. et al. De novo designed proteins neutralize lethal snake venom toxins. Res Sq (2024).

35. Qing, R. et al. Protein Design: From the Aspect of Water Solubility and Stability. Chem Rev 122, 14085–14179 (2022).

36. Jara, J.R., Aroca, P., Solano, F., Martinez, J.H. & Lozano, J.A. The role of sulfhydryl compounds in mammalian melanogenesis: the effect of cysteine and glutathione upon tyrosinase and the intermediates of the pathway. Biochim Biophys Acta 967, 296–303 (1988).

37. Smit, N.P. et al. Melanogenesis in cultured melanocytes can be substantially influenced by L-tyrosine and L-cysteine. J Invest Dermatol 109, 796–800 (1997).

38. Ren, X. et al. The Dominant Role of Oxygen in Modulating the Chemical Evolution Pathways of Tyrosine in Peptides: Dityrosine or Melanin. Angew Chem Int Ed Engl 58, 5872–5876 (2019).

39. Buch, I., Harvey, M.J., Giorgino, T., Anderson, D.P. & De Fabritiis, G. High-throughput all-atom molecular dynamics simulations using distributed computing. J Chem Inf Model 50, 397–403 (2010).

40. Souza, P.C.T. et al. Martini 3: a general purpose force field for coarse-grained molecular dynamics. Nat Methods 18, 382–388 (2021).

41. Wakamatsu, K. & Ito, S. Advanced chemical methods in melanin determination. Pigment Cell Res 15, 174–183 (2002).

42. Camus, A. et al. High conductivity Sepia melanin ink films for environmentally benign printed electronics. Proc Natl Acad Sci U S A 119, e2200058119 (2022).

43. Ito, S. & Wakamatsu, K. Diversity of human hair pigmentation as studied by chemical analysis of eumelanin and pheomelanin. J Eur Acad Dermatol Venereol 25, 1369–1380 (2011).

44. Lamoreux, M.L., Wakamatsu, K. & Ito, S. Interaction of major coat color gene functions in mice as studied by chemical analysis of eumelanin and pheomelanin. Pigment Cell Res 14, 23–31 (2001).

45. Prum, R.O., Torres, R.H., Williamson, S. & Dyck, J. Coherent light scattering by blue feather barbs. Nature 396, 28–29 (1998).

46. Keyser, A.J. & Hill, G.E. Structurally based plumage coloration is an honest signal of quality in male blue grosbeaks. Behavioral Ecology 11, 202–209 (2000).

47. Zi, J. et al. Coloration strategies in peacock feathers. Proc Natl Acad Sci U S A 100, 12576–12578 (2003).

48. Mani, I., Sharma, V., Tamboli, I. & Raman, G. Interaction of melanin with proteins--the importance of an acidic intramelanosomal pH. Pigment Cell Res 14, 170–179 (2001).

49. Canovas, F.G., Garcia-Carmona, F., Sanchez, J.V., Pastor, J.L. & Teruel, J.A. The role of pH in the melanin biosynthesis pathway. J Biol Chem 257, 8738–8744 (1982).

50. Wagh, S., Ramaiah, A., Subramanian, R. & Govindarajan, R. Melanosomal Proteins Promote Melanin Polymerization. Pigment Cell Research 13, 442–448 (2000).

51. MacDonald, J.T. et al. Synthetic beta-solenoid proteins with the fragment-free computational design of a beta-hairpin extension. Proc Natl Acad Sci U S A 113, 10346–10351 (2016).

52. Motovilov, K.A. & Mostert, A.B. Melanin: Nature’s 4th bioorganic polymer. Soft Matter 20, 5635–5651 (2024).

53. Bennett, N.R. et al. Improving de novo protein binder design with deep learning. Nat Commun 14, 2625 (2023).

54. Frederix, P.W. et al. Exploring the sequence space for (tri-)peptide self-assembly to design and discover new hydrogels. Nat Chem 7, 30–37 (2015).

55. de Jong, D.H. et al. Improved Parameters for the Martini Coarse-Grained Protein Force Field. J Chem Theory Comput 9, 687–697 (2013).

56. Hess, B., Kutzner, C., van der Spoel, D. & Lindahl, E. GROMACS 4: Algorithms for Highly Efficient, Load-Balanced, and Scalable Molecular Simulation. J Chem Theory Comput 4, 435–447 (2008).

57. Eisenberg, D., Schwarz, E., Komaromy, M. & Wall, R. Analysis of membrane and surface protein sequences with the hydrophobic moment plot. J Mol Biol 179, 125–142 (1984).

58. Amend, J.P. & Helgeson, H.C. Solubilities of the common L-α-amino acids as a function of temperature and solution pH. Pure & Applied Chemistry 69, 935–942 (1997).

