## Supplementary Information for "Computational design of blue melanin with peptide motif scaffolding"

**Table of Contents:**

**Supplementary Figures**

Supplementary Fig. 1: Motif scaffolding of PRP with RFdiffusion
Supplementary Fig. 2: Experimental characterization of Y-containing pentapeptides
Supplementary Fig. 3: Hybrid blue melanin formed from SPYGG and LAYPS
Supplementary Fig. 4: Alternative techniques to prepare SPYGG blue melanin
Supplementary Fig. 5: Molecular dynamics of SPYGG tight packing
Supplementary Fig. 6: pH stability of synthetic and natural blue pigments
Supplementary Fig. 7: Shelf stability of blue and green melanins
Supplementary Fig. 8: Light and thermal stabilities of SPYGG at neutral and acidic conditions
Supplementary Fig. 9: Light stability of synthetic and natural blue pigments
Supplementary Fig. 10: Thermal stability of synthetic and natural blue pigments

**Supplementary Tables**

Supplementary Table 1: Top 100 Y-containing pentapeptides ranked by solubility
Supplementary Table 2: All 2Y-containing pentapeptides ranked by solubility

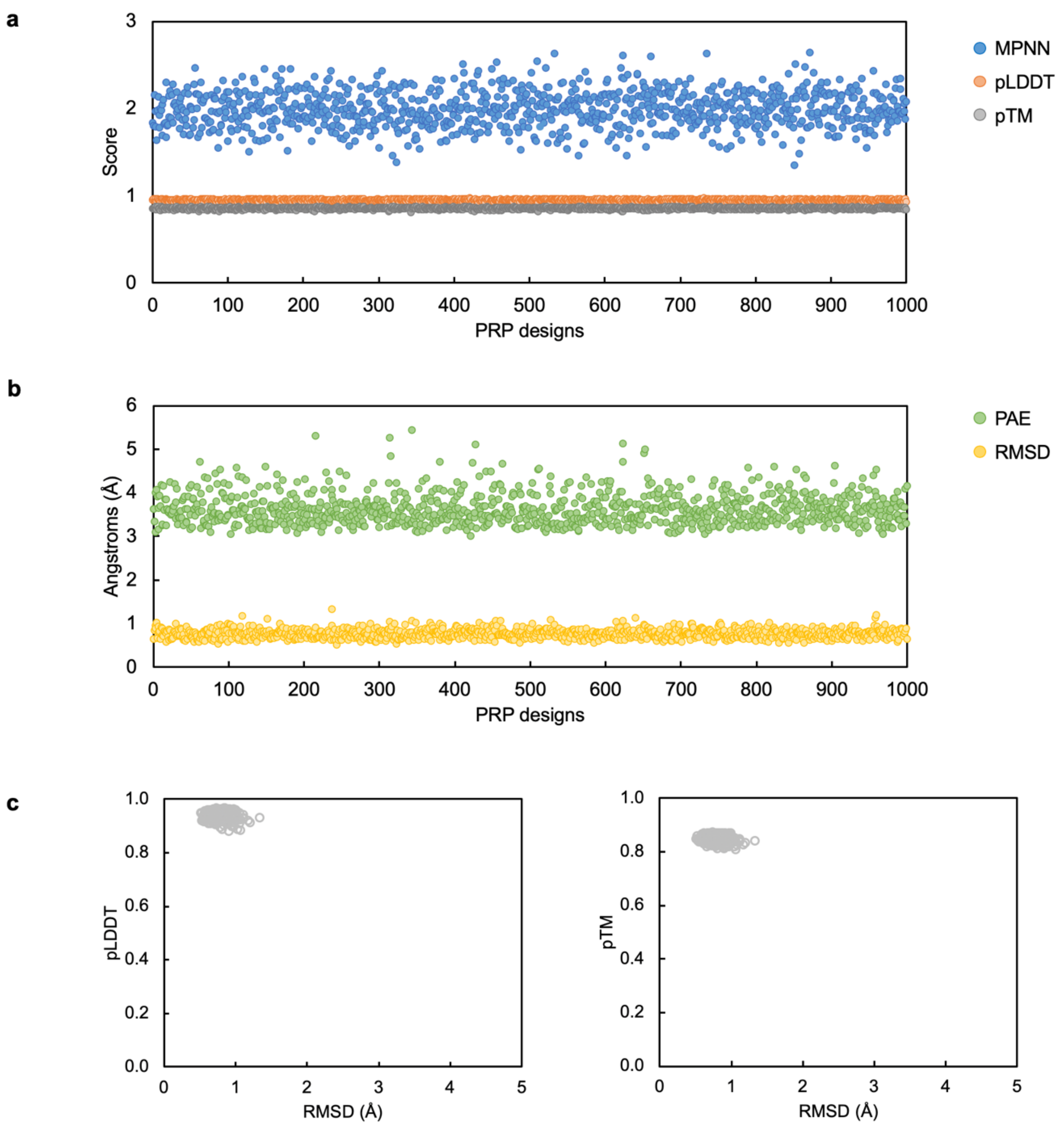


**Supplementary Fig. 1: Motif scaffolding of PRP with RFdiffusion. a**, MPNN, pLDDT, and pTM scores of all 1,000 motif-scaffolded PRPs. **b**, PAE and RMSD scores of all 1,000 motif-scaffolded PRPs. **c**, pLDDT and pTM scores plotted against RMSD.

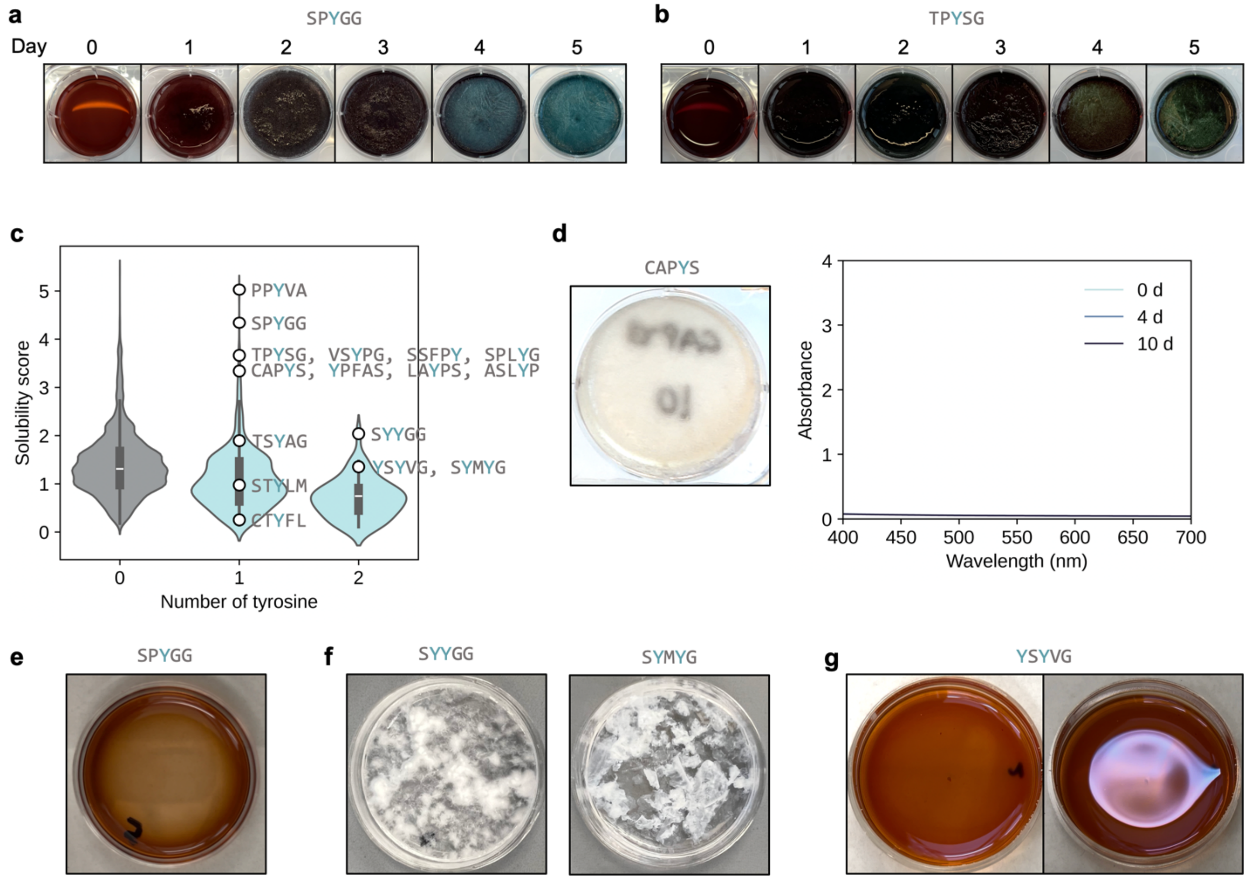


**Supplementary Fig. 2: Experimental characterization of Y-containing pentapeptides. a**, Spontaneous polymerization of SPYGG blue melanin at -20°C for 5 days. **b**, Spontaneous polymerization of TPYSG green melanin at -20°C for 5 days. **c**, Solubility ranking of all experimentally tested peptides. **d**, Absorption spectra of CAPYS melanin at days 0, 4, and 10. **e**, Brown SPYGG melanin after 4 days of polymerization at room temperature. **f**, Aggregation formed from 2Y-containing pentapeptides. **g**, Red-brown YSYVG melanin with an iridescent thin film.

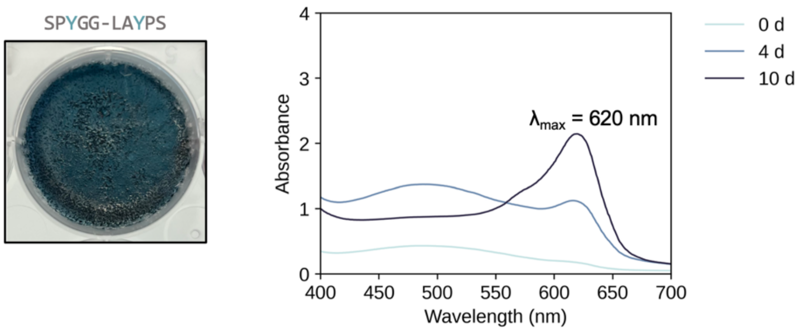


**Supplementary Fig. 3: Hybrid blue melanin formed from SPYGG and LAYPS.** Absorption spectra of blue melanin formed from SPYGG and LAYPS peptides at equal weights (1:1, w/w) at days 0, 4, and 10.

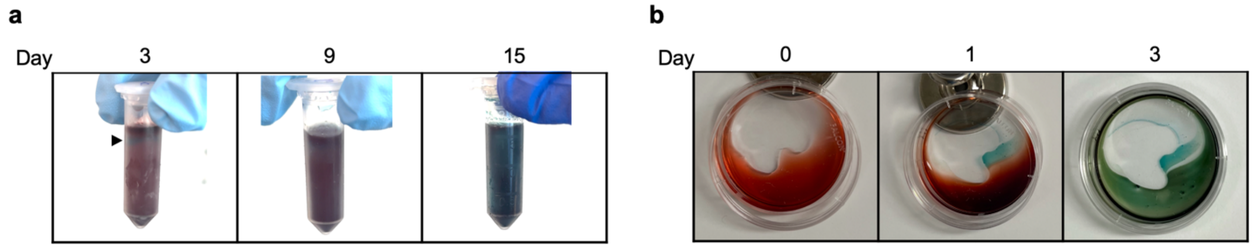


**Supplementary Fig. 4: Alternative techniques to prepare SPYGG blue melanin. a**, Spontaneous polymerization of SPYGG blue melanin at -20°C for 15 days in a 2 mL microcentrifuge tube. Black arrow indicates first blue melanin appearance. **b**, Evaporative polymerization of SPYGG blue melanin at room temperature.

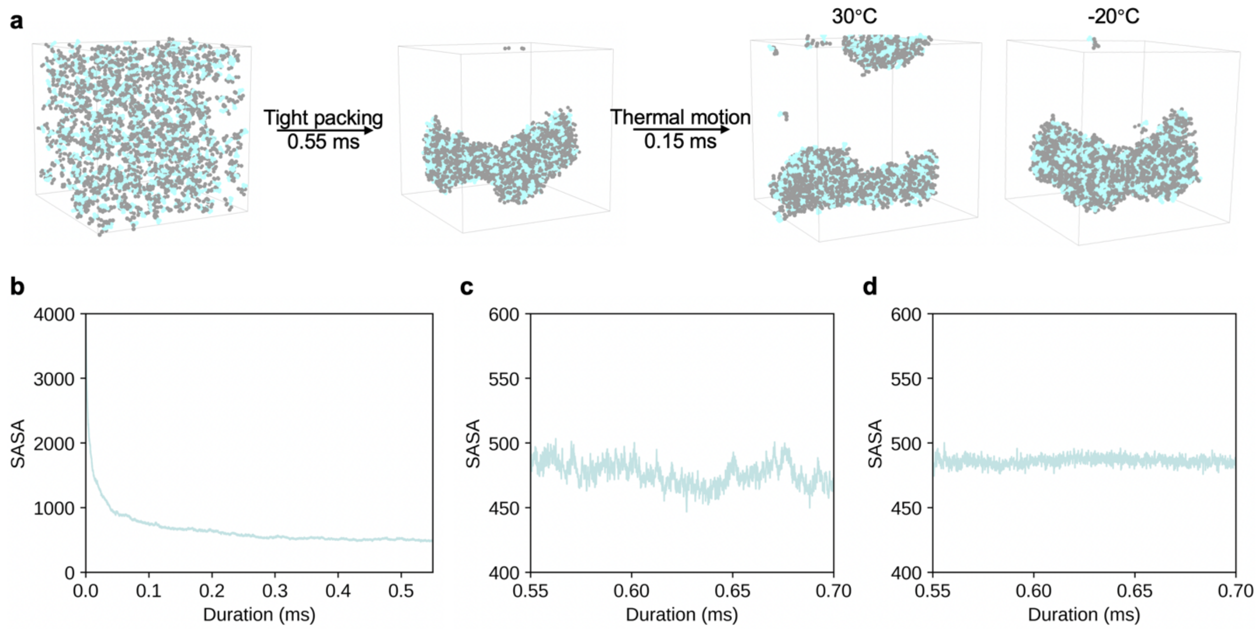


**Supplementary Fig. 5: Molecular dynamics of SPYGG tight packing. a**, Molecular dynamics models of the peptides in water at the beginning and after 0.55 ms of simulation. The peptide model was run for another 0.15 ms at 30°C and -20°C. **b**, Change in SASA in the first 0.55 ms of simulation. Fluctuation in SASA at **c**, 30°C and **d**, -20°C of the tightly packed peptide cluster from 0.55 to 0.70 ms.

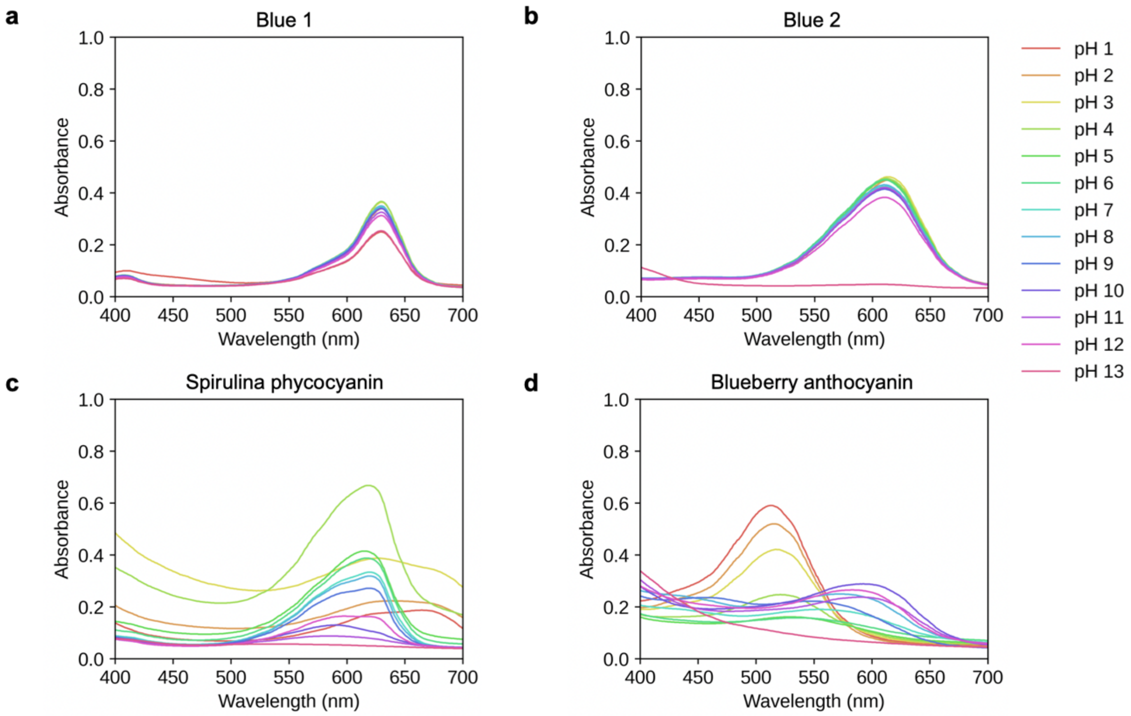


**Supplementary Fig. 6: pH stability of synthetic and natural blue pigments.** Absorption spectra of **a**, FD&C Blue No. 1, **b**, FD&C Blue No. 2, **c**, spirulina phycocyanin, and **d**, blueberry anthocyanin re-suspended in pH 1-13 buffer solutions.

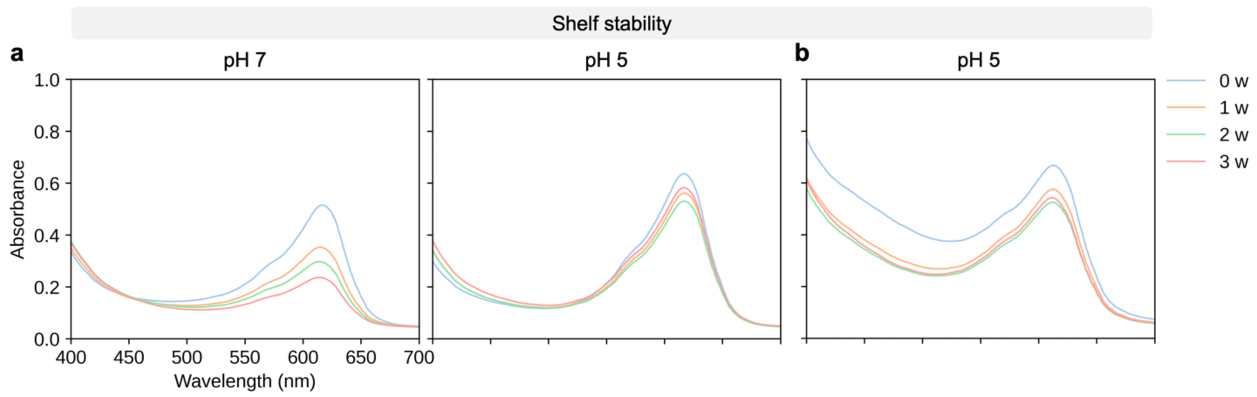


**Supplementary Fig. 7: Shelf stability of blue and green melanins. a**, Absorption spectra of SPYGG blue melanin at pH 7 and 5 kept in the dark at room temperature for 3 weeks. Each spectrum is average of three replicates. **b**, Absorption spectra of TPYSG green melanin re-suspended in pH 5 buffer solution kept in the dark at room temperature for 3 weeks.

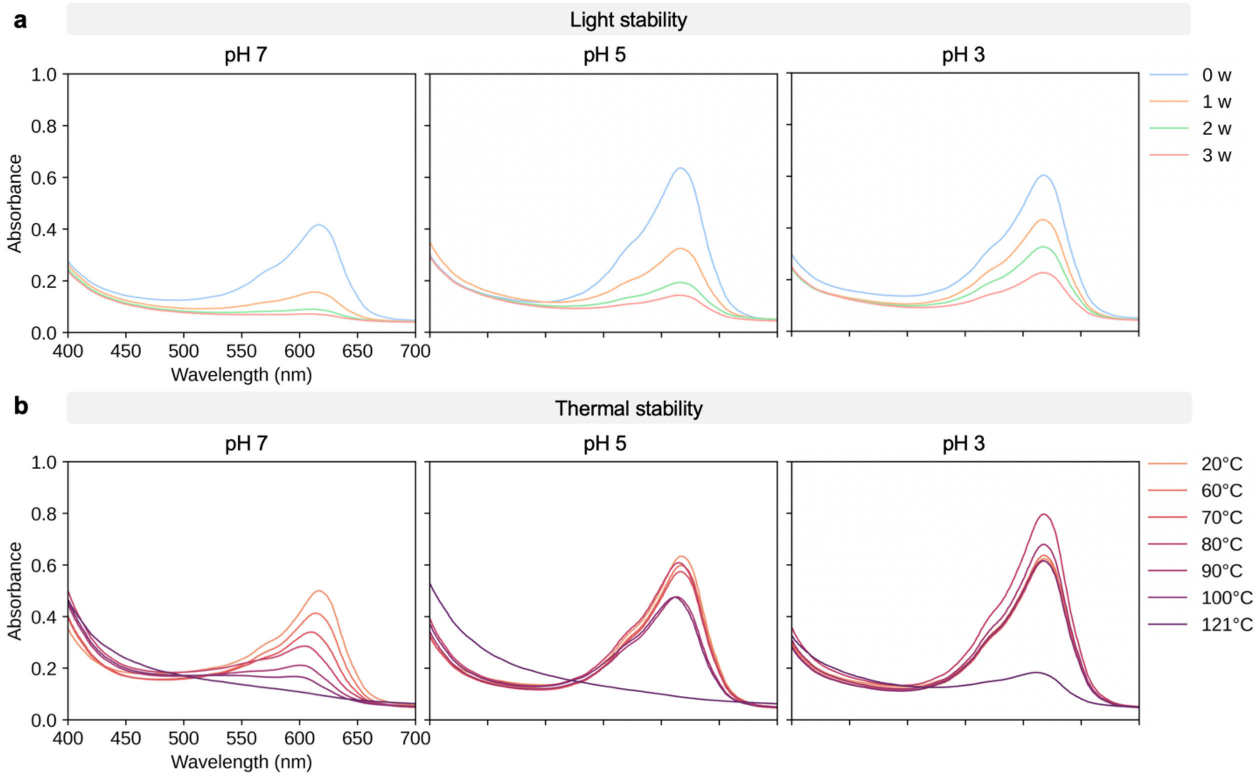


**Supplementary Fig. 8: Light and thermal stabilities of SPYGG at neutral and acidic conditions. a**, Absorption spectra of SPYGG blue melanin at pH 7, 5, and 3 exposed to 600 lux of illumination over 3 weeks. Each spectrum is average of three replicates. **b**, Absorption spectra of SPYGG blue melanin at pH 7, 5, and 3 heated to 60°C, 70°C, 80°C, 90°C, 100°C and 121°C (autoclave). Each spectrum is average of three replicates.

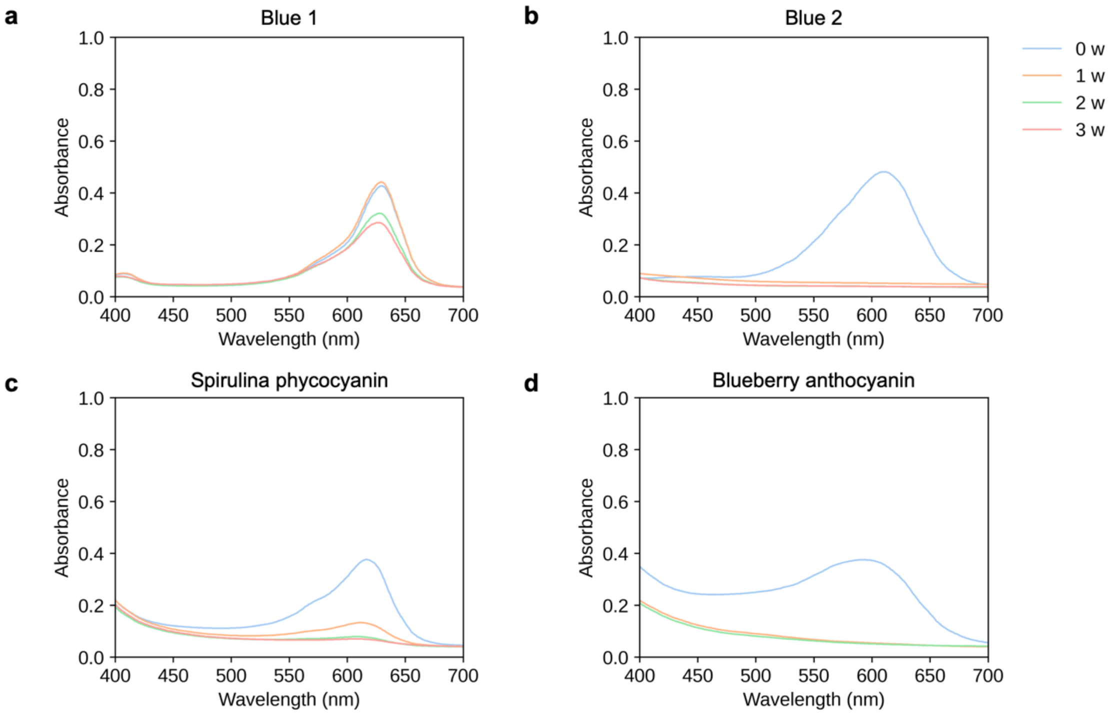


**Supplementary Fig. 9: Light stability of synthetic and natural blue pigments.** Absorption spectra of **a**, FD&C Blue No. 1, **b**, FD&C Blue No. 2, **c**, spirulina phycocyanin, and **d**, blueberry anthocyanin (pH 11) exposed to 600 lux of illumination over 3 weeks.

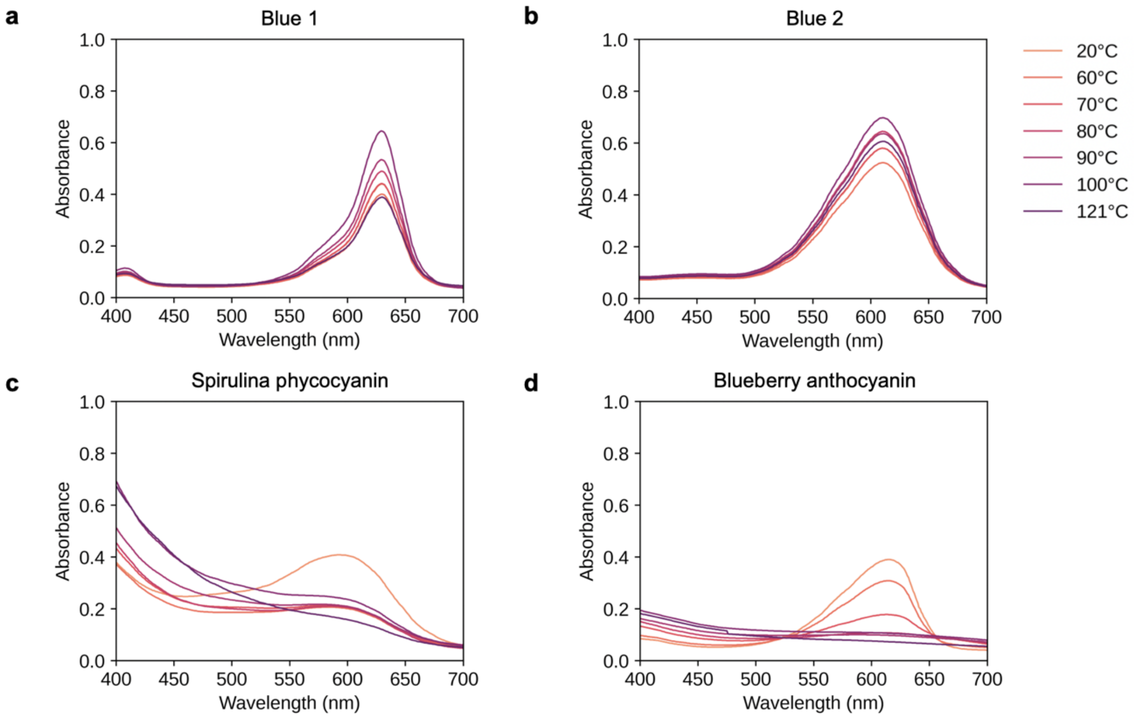


**Supplementary Fig. 10: Thermal stability of synthetic and natural blue pigments.** Absorption spectra of **a**, FD&C Blue No. 1, **b**, FD&C Blue No. 2, **c**, spirulina phycocyanin, and **d**, blueberry anthocyanin (pH 11) heated to 60°C, 70°C, 80°C, 90°C, 100°C and 121°C (autoclave).

**Supplementary Table 1. Top 100 Y-containing pentapeptides ranked by solubility.**

| Ranking | Sequence | Solubility score | Hydrophobicity score |
|---|---|---|---|
| 1 | PPYVA | 5.0 | 2.2 |
| 2 | **SPYGG** | 4.3 | 1.2 |
| 3 | TPYSG | 3.8 | 0.6 |
| 4 | VSYPG | 3.7 | 1.8 |
| 5 | SSFPY | 3.7 | 1.2 |
| 6 | SPLYG | 3.7 | 1.7 |
| 7 | CAPYS | 3.4 | 1.1 |
| 8 | YPFAS | 3.4 | 2.0 |
| 9 | **LAYPS** | 3.4 | 1.9 |
| 10 | ASLYP | 3.4 | 1.9 |
| 11 | STFYP | 3.2 | 1.3 |
| 12 | PTFGY | 3.1 | 2.0 |
| 13 | SVLYP | 3.1 | 2.3 |
| 14 | SVLPY | 3.1 | 2.3 |
| 15 | PSLIY | 3.0 | 2.6 |
| 16 | YLSFP | 3.0 | 2.5 |
| 17 | ATYVP | 2.9 | 2.0 |
| 18 | APTVY | 2.9 | 2.0 |
| 19 | APYVT | 2.9 | 2.0 |
| 20 | TPYAM | 2.9 | 1.6 |
| 21 | AVFPY | 2.8 | 3.3 |
| 22 | AVLYP | 2.8 | 3.1 |
| 23 | CALYP | 2.7 | 2.4 |
| 24 | APLLY | 2.7 | 3.1 |
| 25 | TPYIT | 2.6 | 1.7 |
| 26 | PTLTY | 2.6 | 1.3 |
| 27 | TLPIY | 2.5 | 2.8 |
| 28 | CTYPF | 2.5 | 1.8 |
| 29 | VPFVY | 2.5 | 3.7 |
| 30 | TLLPY | 2.5 | 2.5 |
| 31 | VPIYL | 2.4 | 3.9 |
| 32 | ASYSG | 2.4 | 1.0 |
| 33 | CVFPY | 2.4 | 2.9 |
| 34 | VLFPY | 2.4 | 3.7 |
| 35 | PVFLY | 2.4 | 3.7 |
| 36 | PILIY | 2.4 | 4.2 |
| 37 | IPCYF | 2.4 | 3.2 |
| 38 | CYLIP | 2.4 | 3.1 |
| 39 | CLLPY | 2.4 | 2.8 |
| 40 | PLFLY | 2.4 | 3.7 |
| 41 | PLLYF | 2.4 | 3.7 |
| 42 | STYSG | 2.2 | 0.3 |
| 43 | SSYTG | 2.2 | 0.3 |
| 44 | STYSG | 2.2 | 0.3 |
| 45 | SSYTG | 2.2 | 0.3 |
| 46 | SSYVG | 2.2 | 1.5 |
| 47 | YSFSS | 2.1 | 0.9 |
| 48 | SSFYS | 2.1 | 0.9 |
| 49 | AYAGS | 2.1 | 1.8 |
| 50 | SSCYG | 2.1 | 0.7 |

| | | | |
|---|---|---|---|
| 51 | YSFSG | 2.1 | 1.6 |
| 52 | SYFSG | 2.1 | 1.6 |
| 53 | SYFSG | 2.1 | 1.6 |
| 54 | SSFYG | 2.1 | 1.6 |
| 55 | SSLGY | 2.1 | 1.4 |
| 56 | SGLYG | 2.1 | 2.1 |
| 57 | GYFGG | 2.1 | 2.9 |
| 58 | TSAGY | 1.9 | 1.1 |
| 59 | TSYAG | 1.9 | 1.1 |
| 60 | TYGGA | 1.9 | 1.8 |
| 61 | SVYSA | 1.9 | 1.6 |
| 62 | YSAVS | 1.9 | 1.6 |
| 63 | AVYSG | 1.8 | 2.3 |
| 64 | ASYVG | 1.8 | 2.3 |
| 65 | AVYSG | 1.8 | 2.3 |
| 66 | AYMSS | 1.8 | 1.2 |
| 67 | SSFYA | 1.8 | 1.7 |
| 68 | SYFAS | 1.8 | 1.7 |
| 69 | SSFYA | 1.8 | 1.7 |
| 70 | SSFYA | 1.8 | 1.7 |
| 71 | SSFAY | 1.8 | 1.7 |
| 72 | SSYAL | 1.8 | 1.6 |
| 73 | SYLSA | 1.8 | 1.6 |
| 74 | AYLSS | 1.8 | 1.6 |
| 75 | SLASY | 1.8 | 1.6 |
| 76 | SSLYA | 1.8 | 1.6 |
| 77 | ASLSY | 1.8 | 1.6 |
| 78 | ASLSY | 1.8 | 1.6 |
| 79 | SSLAY | 1.8 | 1.6 |
| 80 | AYLSS | 1.8 | 1.6 |
| 81 | ASIYG | 1.8 | 2.6 |
| 82 | ISGYA | 1.8 | 2.6 |
| 83 | SAIYG | 1.8 | 2.6 |
| 84 | AYIGS | 1.8 | 2.6 |
| 85 | CAYSG | 1.8 | 1.5 |
| 86 | ASFYG | 1.8 | 2.4 |
| 87 | ASFYG | 1.8 | 2.4 |
| 88 | SAFYG | 1.8 | 2.4 |
| 89 | ASFYG | 1.8 | 2.4 |
| 90 | GSFYA | 1.8 | 2.4 |
| 91 | ASFYG | 1.8 | 2.4 |
| 92 | AGFYS | 1.8 | 2.4 |
| 93 | ALYSG | 1.8 | 2.2 |
| 94 | LYASG | 1.8 | 2.2 |
| 95 | SALYG | 1.8 | 2.2 |
| 96 | SYLAG | 1.8 | 2.2 |
| 97 | ASLYG | 1.8 | 2.2 |
| 98 | AGFYG | 1.8 | 3.0 |
| 99 | STYTS | 1.7 | -0.2 |
| 100 | SSVTY | 1.7 | 0.9 |

**Bolded.** Successful blue melanin formation.

**Supplementary Table 2. All 2Y-containing pentapeptides ranked by solubility.**

| Ranking | Sequence | Solubility score | Hydrophobicity score |
|---|---|---|---|
| 1 | SYYGG | 2.0 | 1.3 |
| 2 | YSYVG | 1.5 | 1.9 |
| 3 | SYMYG | 1.4 | 1.5 |
| 4 | YSFYG | 1.4 | 2.0 |
| 5 | SGLYY | 1.4 | 1.9 |
| 6 | ASFYY | 1.1 | 2.2 |
| 7 | SYFAY | 1.1 | 2.2 |
| 8 | AFYYS | 1.1 | 2.2 |
| 9 | ASLYY | 1.1 | 2.0 |
| 10 | AGLYY | 1.1 | 2.7 |
| 11 | TSYYV | 1.0 | 1.4 |
| 12 | STMYY | 0.9 | 0.9 |
| 13 | TYMYG | 0.9 | 1.6 |
| 14 | YYLTS | 0.9 | 1.4 |
| 15 | GTFYY | 0.9 | 2.1 |
| 16 | SVFYY | 0.8 | 2.6 |
| 17 | AFYYA | 0.8 | 3.0 |
| 18 | YSFIY | 0.8 | 2.9 |
| 19 | YFCYS | 0.8 | 1.8 |
| 20 | SFLYY | 0.8 | 2.6 |
| 21 | SLFYY | 0.8 | 2.6 |
| 22 | IYFGY | 0.8 | 3.6 |
| 23 | CYFYG | 0.8 | 2.5 |
| 24 | CGYYF | 0.8 | 2.5 |
| 25 | LGFYY | 0.7 | 3.3 |
| 26 | GLFYY | 0.7 | 3.3 |
| 27 | GLYLY | 0.7 | 3.1 |
| 28 | CTYYA | 0.6 | 1.4 |
| 29 | TAFYY | 0.6 | 2.3 |
| 30 | ATYYL | 0.6 | 2.2 |
| 31 | AIVYY | 0.5 | 3.6 |
| 32 | AYFYV | 0.5 | 3.4 |
| 33 | AVYLY | 0.5 | 3.3 |
| 34 | AYFIY | 0.5 | 3.7 |
| 35 | CYFYA | 0.5 | 2.6 |
| 36 | CAFYY | 0.5 | 2.6 |
| 37 | CAFYY | 0.5 | 2.6 |
| 38 | AYLYF | 0.4 | 3.4 |
| 39 | AFLYY | 0.4 | 3.4 |
| 40 | ALLYY | 0.4 | 3.3 |
| 41 | VYFYT | 0.3 | 2.7 |
| 42 | YTFFY | 0.2 | 2.8 |
| 43 | CVFYY | 0.2 | 3.1 |
| 44 | CVFYY | 0.2 | 3.1 |
| 45 | CVFYY | 0.2 | 3.1 |
| 46 | CVLYY | 0.2 | 3.0 |
| 47 | YFFIY | 0.1 | 4.3 |
| 48 | YYLIL | 0.1 | 4.0 |
| 49 | CLFYY | 0.1 | 3.1 |
| 50 | LFFYY | 0.1 | 4.0 |
